# CHARTS: A web application for characterizing and comparing tumor subpopulations in publicly available single-cell RNA-seq datasets

**DOI:** 10.1101/2020.09.23.310441

**Authors:** Matthew N. Bernstein, Zijian Ni, Michael Collins, Mark E. Burkard, Christina Kendziorski, Ron Stewart

## Abstract

**Background:** Single-cell RNA-seq (scRNA-seq) enables the profiling of genome-wide gene expression at the single-cell level and in so doing facilitates insight into and information about cellular heterogeneity within a tissue. Perhaps nowhere is this more important than in cancer, where tumor and tumor microenvironment heterogeneity directly impact development, maintenance, and progression of disease. While publicly available scRNA-seq cancer datasets offer unprecedented opportunity to better understand the mechanisms underlying tumor progression, metastasis, drug resistance, and immune evasion, much of the available information has been underutilized, in part, due to the lack of tools available for aggregating and analysing these data.

**Results:** We present CHARacterizing Tumor Subpopulations (CHARTS), a computational pipeline and web application for analyzing, characterizing, and integrating publicly available scRNA-seq cancer datasets. CHARTS enables the exploration of individual gene expression, cell type, malignancy-status, differentially expressed genes, and gene set enrichment results in subpopulations of cells across multiple tumors and datasets.

**Conclusion:** CHARTS is an easy to use, comprehensive platform for exploring single-cell subpopulations within tumors across the ever-growing collection of public scRNA-seq cancer datasets. CHARTS is freely available at charts.morgridge.org.

## Introduction

Over the past three decades, the cancer research community has amassed large quantities of gene expression data from tumors. The premier example of such data was generated by The Cancer Genome Atlas (Cancer Genome Atlas Research Network et al., 2013), which generated bulk RNA-seq and microarray data from thousands of tumors across dozens of cancer types. These data have enabled a greater understanding into the molecular biology of cancer and have revealed great heterogeneity not only between cancer types, but also between tumors of the same cancer type (Bedard et al., 2013). Unfortunately, investigations utilizing this resource are limited by the fact that gene expression was profiled using bulk methods, which measure gene expression on average across thousands, or tens of thousands, of cells in a sample. With the advent of single-cell RNA-seq (scRNA-seq), investigators are now able to measure gene expression at the single-cell level thereby gaining access to the substantial heterogeneity of cells within a tumor and the tumor microenvironment (González-Silva et al., 2020). Publicly available scRNA-seq cancer datasets offer unprecedented opportunity to better understand the mechanisms of tumor progression, metastasis, drug resistance, and immune evasion. However, analyzing these data in the aggregate is challenging, especially for those without strong computational skills. To this end, easy-to-use web-based tools are important for enabling the broader research community to perform integrative analyses and, in doing so, to increase their ability to leverage their knowledge and comprehensively examine scientific and/or clinically relevant hypotheses in multiple datasets.

While a few web-based tools for analyzing scRNA-seq data are available, they are not designed specifically for cancer research or do not easily enable exploration of existing public datasets. For example, recent tools such as Alona (Franzén and Björkegren, 2020) and Granatum (Zhu et al., 2017) enable scRNA-seq analysis in the web browser; however, these tools are not cancer-specific and therefore do not enable important cancer-specific tasks such as classifying cells as being either transformed malignant cells or untransformed cells of the tumor microenvironment. Furthermore, these tools do not enable exploration of preprocessed, publicly available scRNA-seq datasets. Another tool, GREIN (Mahi et al., 2019), enables exploration of public gene expression data, but it is neither single-cell specific nor cancer-specific and, consequently, does not implement features necessary for single-cell analysis such as cell type identification, clustering, or gene set enrichment, nor does it implement cancer-specific analyses such as malignancy classification. CancerSEA (Yuan et al., 2019) enables exploration of gene set enrichment scores for gene sets pertaining to cancer-related processes, but does not enable visualization, differential expression, or cell type identification. In short, while web-based tools exist for exploring expression data, most do not allow for detailed analysis of scRNA-seq data across diverse tumors and datasets.

To address this gap, we present CHARacterizing Tumor Subpopulations (CHARTS), a web application and associated computational pipeline for analyzing and characterizing cancer scRNA-seq datasets. As described in detail below, for each tumor in its database, CHARTS identifies clusters and enables exploration via interactive dimension-reduction methods. Derived clusters are annotated with cell types from the Cell Ontology (Bard et al., 2005) via CellO (Bernstein et al., 2020), with information provided on the probability of the specific cell type as well as its ancestors. For example, the data may provide substantial evidence to classify cells within a cluster as T cells, but less evidence may be available to classify cells into more specific functional groups (e.g. helper or memory T cells). In addition, for each cluster within each tumor, enrichment of genes involved in biological processes and pathways is provided. Genes that are differentially expressed between the cluster and others are also available. Finally, CHARTS can be used to distinguish malignant vs. non-malignant cells allowing for precise exploration into the interactions between cell subpopulations within the tumor microenvironment. CHARTS currently enables exploration of 61 tumors across six cancer types, and data is being continually added. CHARTS is freely available at charts.morgridge.org.

## Implementation

Publicly available expression data were downloaded from the Gene Expression Omnibus (Edgar et al., 2002) and normalized to units of log(TPM+1). An offline computational pipeline implements a number of analyses in order to enable comprehensive characterization and comparison of tumor subpopulations within and between tumors (Fig. 1). All analyses output is stored in a backend database, which is quickly and easily accessible to a user through a frontend web application.

**Figure 1.**
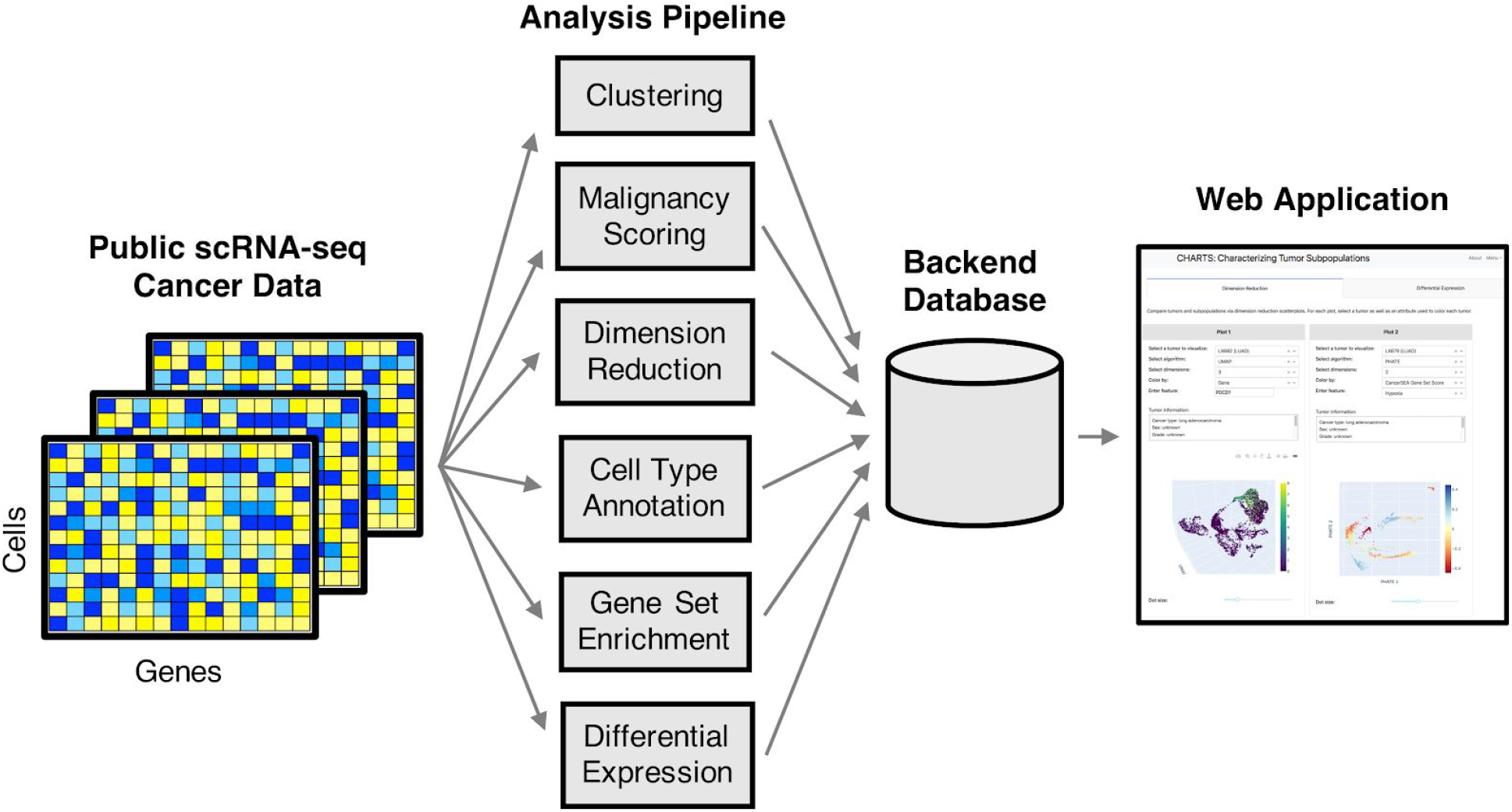
Overview. A schematic diagram of the CHARTS pipeline. Public scRNA-seq datasets are collected and analyzed with a custom pipeline. This pipeline computes clusters, malignancy scores, dimension reduction transformations, cell type annotations, gene set enrichment scores, and differentially expressed genes for each cluster. Results are stored in a backend database and are accessed from the frontend web application.

### Dimension Reduction

A user may construct interactive dimension-reduction scatterplots using two or three-dimensional UMAP (McInnes et al.) or PHATE (Moon et al., 2019). Each cell can be colored by the expression of a user-specified gene, cluster, malignancy score, cell type, or gene set enrichment. Two scatterplots are placed side-by-side enabling users to compare two characteristics (e.g. two different genes’ expression values or a gene’s expression value and the predicted cell types) within the same tumor or to compare a single characteristic (e.g. a single gene’s expression values) between two tumors.

### Clustering

Clustering is performed using the Leiden community detection algorithm (Traag et al., 2019), as implemented in Python’s Scanpy library (Wolf et al., 2018). Leiden’s resolution parameter is set to 4.0, a higher value than the default resolution of 1.0, due to the empirical observation that for tumor datasets consisting of thousands of cells, the default resolution of 1.0 resulted in clusters that combined multiple putative cell types (data not shown).

### Cell Type Annotation

Each cluster is annotated with cell types from the Cell Ontology (Bard et al., 2005) via CellO (Bernstein et al., 2020). The Cell Ontology is a hierarchically structured knowledgebase of known cell types. Specifically, the Cell Ontology forms a directed acyclic graph (DAG) where edges in the graph represent “is a” relationships. Because of this DAG structure, each cell is assigned to a specific cell type as well as all ancestors of this specific cell type within the DAG. CellO was trained using the isotonic regression correction algorithm. CHARTS exposes both CellO’s binary cell type decisions for each cell type as well as CellO’s estimated probability that each cell is of a given type.

### Gene Set Enrichment

Each cluster’s mean gene expression profile is scored for enrichment of gene sets describing molecular processes. Specifically, CHARTS uses GSVA (Hänzelmann et al., 2013) to score each cluster for enrichment of gene sets in the hallmark gene set collection from MSigDB (Liberzon et al., 2015) and the gene set collection used by CancerSEA.

### Malignancy Status

Each cell is assigned a malignancy score that describes the likelihood that the cell is malignant. The malignancy scoring approach builds upon the approaches used by (Tirosh et al., 2016) and (Couturier et al., 2020) for classifying cells as either transformed, malignant cells or untransformed cells within the tumor microenvironment (Supplementary Methods).

### Differential Expression

For each cluster within each tumor, CHARTS uses a Wilcoxon rank-sum test, as implemented in Scanpy, to compute the set of genes differentially expressed in the given cluster versus cells outside the cluster within the given tumor. CHARTS presents all genes meeting a false discovery rate threshold of 0.05 after correction via the Benjamini-Hochberg procedure (Benjamini and Hochberg, 1995).

### Implementation Details

The CHARTS website is implemented with Plotly’s Dash framework (https://plotly.com/dash/). The data analysis pipeline was implemented with Snakemake (Köster and Rahmann, 2012). The software that implements the website and backend pipeline is freely available for users to run CHARTS locally in order to explore their own data alongside existing public data.

## Results

Two case studies demonstrate how CHARTS can be used, both to examine and generate new hypotheses.

### Case study: dysfunctional CD8+ T cells in lung adenocarcinoma

Investigators have recently reported a dysfunctional population of CD8+ T cells in lung cancer (Thommen et al., 2018) and melanoma (Li et al., 2019) that express genes associated with immune suppression. In some melanoma samples, this population was also found to be highly proliferative (Li et al., 2019). We used CHARTS to explore whether this dysfunctional state was common across the majority of CD8+ T cells, and to evaluate whether dysfunctional CD8+ T cells were also highly proliferative. We found that in the majority of lung adenocarcinomas, only a subset of CD8+T cells express marker genes for this dysfunctional state. Two adenocarcinomas from Laughney et al. (2020) are shown in Fig. 2. Using the gene set enrichment feature of CHARTS, we further found that dysfunctional cells are enriched for cell cycle genes, which may indicate that these dysfunctional CD8+ T cells are highly proliferative in lung adenocarcinoma, as has been recently observed in melanoma.

**Figure 2.**
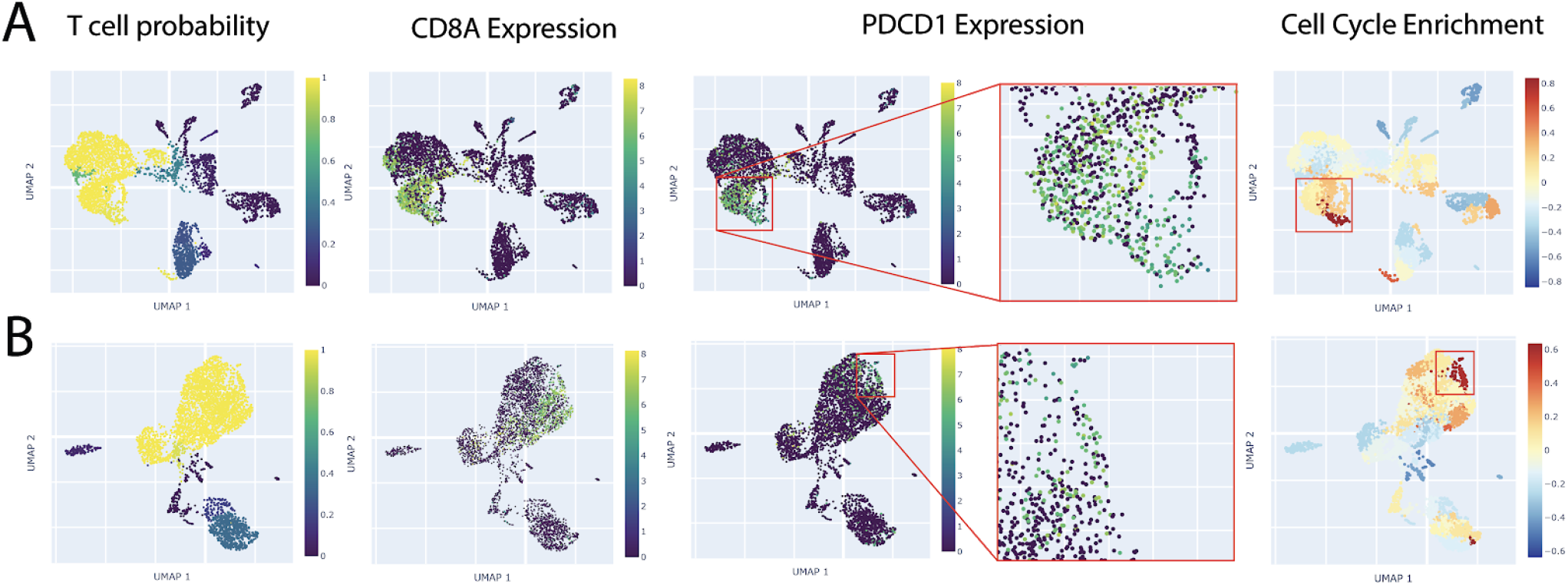
Dysfunctional CD8+ T Cells in Lung Adenocarcinoma. For lung adenocarcinoma tumors LX676 (**A**) and LX682 (**B**), we used CHARTS to visualize the probability that each cell is a T cell as well as expression of CD8A, expression of the dysfunctional CD8+ T cell marker PDCD1, and each cell’s enrichment score for genes in CancerSEA’s cell cycle gene set as produced by GSVA.

### Case study: monocarboxylate transporters in glioblastoma

We investigated the expression of MCT4, a prognostic biomarker of glioblastoma aggression (Lai et al., 2020; Zuo et al., 2019). Using CHARTS, we found that MCT4 tended to be expressed in the myeloid tumor-infiltrating immune cells. Two tumors from Yuan et al. (2018) are shown in Fig. 3A and 3B. While MCT4 is known to be involved in a metabolic symbiosis between hypoxic tumor cells, which express MCT4 to expel lactate, and oxidative tumor cells, which express MCT1 to intake lactate (Payen et al., 2020) (Fig.3C), the specific cell types expressing MCT4 in glioblastoma have not been well characterized. We used CHARTS to determine which cells express MCT1 in glioblastoma and found that this gene was primarily expressed in cells with high malignancy scores (Fig. 3A, B). Using the gene set enrichment feature of CHARTS, we observed that cells expressing MCT1 tended to express genes enriched for hypoxia, whereas cells expressing MCT4 tended to express genes that were less enriched for hypoxia (Fig. 3A, B). This observation indicates a possible metabolic symbiosis between malignant cells and myeloid cells in the tumor microenvironment of glioblastoma, which to the best of our knowledge, has not been well characterized.

**Figure 3.**
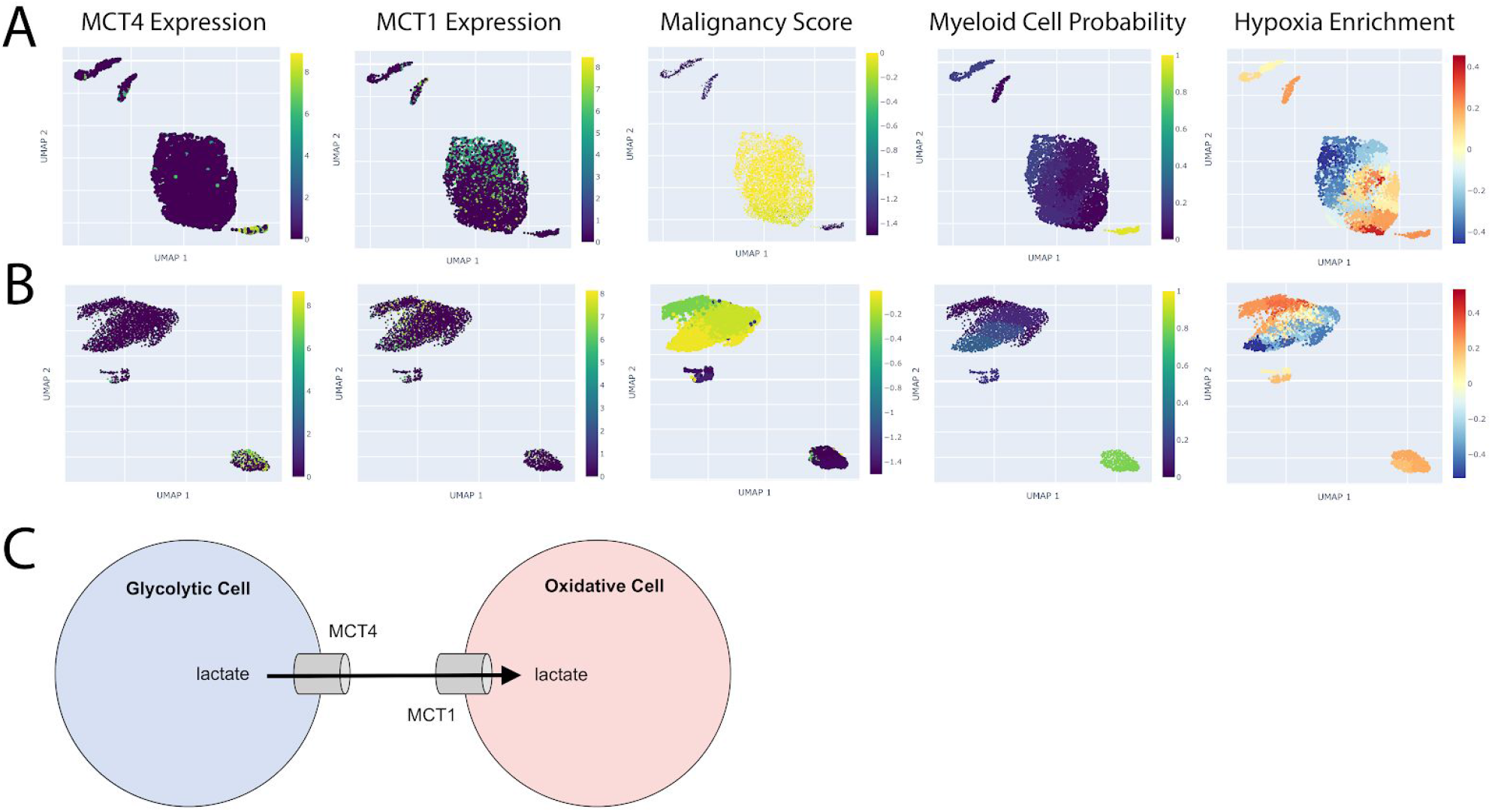
Monocarboxylate Transporter Expression in Glioblastoma. For glioblastoma tumors PJ025 (**A**) and PJ035 (**B**), we used CHARTS to visualize the expression of MCT4, the expression of MCT1, malignancy score, the probability that each cell is a myeloid cell, and each cell’s enrichment score for genes in the Hallmark hypoxia gene set as produced by GSVA. (**C**) A schematic illustration of the metabolic symbiosis between hypoxic, glycolytic tumor cells expressing MCT4 and oxidative tumor cells expressing MCT1.

## Conclusion

In this work, we present CHARTS: a comprehensive framework for exploring single-cell subpopulations within tumors and the tumor microenvironment across ever-growing datasets. CHARTS can be used to develop and explore new hypotheses underlying tumor progression, drug resistance, and immune evasion.

## Availability of data and materials

Code implementing the web application and offline data analysis pipeline is available at https://github.com/stewart-lab/CHARTS.

## Authors’ contributions

M.N.B. lead the conceptual design and implementation of the project and also wrote the manuscript. M.N.B. and Z.N. developed the malignancy scoring technique. M.C. provided technical support and set up the web server. M.E.B. provided input as well as biological and medical expertise. C.K. and R.S. supervised the project.

## References

Bard, J., Rhee, S.Y., and Ashburner, M. (2005). 10.1186/gb-2005-6-2-r21. Genome Biol 6, R21.

Bedard, P.L., Hansen, A.R., Ratain, M.J., and Siu, L.L. (2013). Tumour heterogeneity in the clinic. Nature 501, 355–364.

Benjamini, Y., and Hochberg, Y. (1995). Controlling the False Discovery Rate: A Practical and Powerful Approach to Multiple Testing. Journal of the Royal Statistical Society: Series B (Methodological) 57, 289–300.

Bernstein, M.N., Ma, Z., Gleicher, M., and Dewey, C.N. (2020). CellO: Comprehensive and hierarchical cell type classification of human cells with the Cell Ontology.

Cancer Genome Atlas Research Network, Weinstein, J.N., Collisson, E.A., Mills, G.B., Shaw, K.R.M., Ozenberger, B.A., Ellrott, K., Shmulevich, I., Sander, C., and Stuart, J.M. (2013). The Cancer Genome Atlas Pan-Cancer analysis project. Nat. Genet. 45, 1113–1120.

Couturier, C.P., Ayyadhury, S., Le, P.U., Nadaf, J., Monlong, J., Riva, G., Allache, R., Baig, S., Yan, X., Bourgey, M., et al. (2020). Single-cell RNA-seq reveals that glioblastoma recapitulates a normal neurodevelopmental hierarchy. Nat. Commun. 11, 3406.

Edgar, R., Domrachev, M., and Lash, A.E. (2002). Gene Expression Omnibus: NCBI gene expression and hybridization array data repository. Nucleic Acids Res. 30, 207–210.

Franzén, O., and Björkegren, J.L.M. (2020). alona: a web server for single-cell RNA-seq analysis. Bioinformatics 36, 3910–3912.

González-Silva, L., Quevedo, L., and Varela, I. (2020). Tumor Functional Heterogeneity Unraveled by scRNA-seq Technologies. Trends Cancer Res. 6, 13–19.

Hänzelmann, S., Castelo, R., and Guinney, J. (2013). GSVA: gene set variation analysis for microarray and RNA-Seq data. BMC Bioinformatics 14, 7.

Köster, J., and Rahmann, S. (2012). Snakemake--a scalable bioinformatics workflow engine. Bioinformatics 28, 2520–2522.

Lai, S.-W., Lin, H.-J., Liu, Y.-S., Yang, L.-Y., and Lu, D.-Y. (2020). Monocarboxylate Transporter 4 Regulates Glioblastoma Motility and Monocyte Binding Ability. Cancers 12.

Laughney, A.M., Hu, J., Campbell, N.R., Bakhoum, S.F., Setty, M., Lavallée, V.-P., Xie, Y., Masilionis, I., Carr, A.J., Kottapalli, S., et al. (2020). Regenerative lineages and immune-mediated pruning in lung cancer metastasis. Nat. Med. 26, 259–269.

Li, H., van der Leun, A.M., Yofe, I., Lubling, Y., Gelbard-Solodkin, D., van Akkooi, A.C.J., van den Braber, M., Rozeman, E.A., Haanen, J.B.A.G., Blank, C.U., et al. (2019). Dysfunctional CD8 T Cells Form a Proliferative, Dynamically Regulated Compartment within Human Melanoma. Cell 176, 775–789.e18.

Liberzon, A., Birger, C., Thorvaldsdóttir, H., Ghandi, M., Mesirov, J.P., and Tamayo, P. (2015). The Molecular Signatures Database (MSigDB) hallmark gene set collection. Cell Syst 1, 417–425.

Mahi, N.A., Najafabadi, M.F., Pilarczyk, M., Kouril, M., and Medvedovic, M. (2019). GREIN: An Interactive Web Platform for Re-analyzing GEO RNA-seq Data. Sci. Rep. 9, 7580.

McInnes, L., Healy, J., and Melville, J. UMAP: Uniform Manifold Approximation and Projection for Dimension Reduction. arXiv.

Moon, K.R., van Dijk, D., Wang, Z., Gigante, S., Burkhardt, D.B., Chen, W.S., Yim, K., van den Elzen, A., Hirn, M.J., Coifman, R.R., et al. (2019). Visualizing structure and transitions in high-dimensional biological data. Nat. Biotechnol. 37, 1482–1492.

Payen, V.L., Mina, E., Van Hée, V.F., Porporato, P.E., and Sonveaux, P. (2020). Monocarboxylate transporters in cancer. Mol Metab 33, 48–66.

Thommen, D.S., Koelzer, V.H., Herzig, P., Roller, A., Trefny, M., Dimeloe, S., Kiialainen, A., Hanhart, J., Schill, C., Hess, C., et al. (2018). A transcriptionally and functionally distinct PD-1 CD8 T cell pool with predictive potential in non-small-cell lung cancer treated with PD-1 blockade. Nat. Med. 24, 994–1004.

Tirosh, I., Izar, B., Prakadan, S.M., Wadsworth, M.H., 2nd, Treacy, D., Trombetta, J.J., Rotem, A., Rodman, C., Lian, C., Murphy, G., et al. (2016). Dissecting the multicellular ecosystem of metastatic melanoma by single-cell RNA-seq. Science 352, 189–196.

Traag, V.A., Waltman, L., and van Eck, N.J. (2019). From Louvain to Leiden: guaranteeing well-connected communities. Scientific Reports 9.

Wolf, F.A., Angerer, P., and Theis, F.J. (2018). SCANPY: large-scale single-cell gene expression data analysis. Genome Biol. 19, 15.

Yuan, H., Yan, M., Zhang, G., Liu, W., Deng, C., Liao, G., Xu, L., Luo, T., Yan, H., Long, Z., et al. (2019). CancerSEA: a cancer single-cell state atlas. Nucleic Acids Res. 47, D900–D908.

Yuan, J., Levitin, H.M., Frattini, V., Bush, E.C., Boyett, D.M., Samanamud, J., Ceccarelli, M., Dovas, A., Zanazzi, G., Canoll, P., et al. (2018). Single-cell transcriptome analysis of lineage diversity in high-grade glioma. Genome Med. 10, 57.

Zhu, X., Wolfgruber, T.K., Tasato, A., Arisdakessian, C., Garmire, D.G., and Garmire, L.X. (2017). Granatum: a graphical single-cell RNA-Seq analysis pipeline for genomics scientists. Genome Med. 9, 108.

Zuo, S., Zhang, X., and Wang, L. (2019). A RNA sequencing-based six-gene signature for survival prediction in patients with glioblastoma. Sci. Rep. 9, 2615.

