## Supplemental Methods for "CHARTS: A web application for characterizing and comparing tumor subpopulations in publicly available single-cell RNA-seq datasets"

*Description of Malignancy Score*

We extend the method described by Tirosh et al. (2016) and Couturier et al. (2020) for classifying transformed, malignant cells from untransformed cells. For each cell, we estimate the copy number of each gene by sliding a window of 100 genes down each chromosome and compute the mean expression within each window. These window-mean expression profiles are clustered across all cells *within a given study* (i.e. within the dataset associated with a publication that consists of cells from multiple tumors). The cells within each cluster are then scored according to the tumor-purity of the cluster it belongs to. That is, clusters that consist of cells from multiple tumors will be assigned a lower-malignancy score, whereas tumor clusters that consist of cells from one tumor will be given a high malignancy score. The idea here is that malignant cells within a given tumor will generally have a unique aberrant copy number signature and thus, will cluster only with malignant cells from the same tumor. We devise a quantitative measure of “tumor purity” for a given cluster by computing the entropy associated with the tumor proportions within the tumor. Specifically, we let $p_{i,1}, p_{i,2}, \ldots, p_{i,m}$ be the proportions of cells from tumors $1, \ldots, m$ within cluster $i$ where

$$\sum_{j=1}^{m} p_{i,j} = 1$$

The malignancy score given to cells within cluster $i$ is then the negative entropy of a categorical random variable with probabilities $p_{i,1}, \ldots, p_{i,m}$:

$$\text{Score}_i := \sum_{j=1}^{m} p_{i,j} \log p_{i,j}$$

*Benchmarking Malignancy Score*

We assessed the malignancy score using melanoma data from (Tirosh et al., 2016) (Fig. 1A-D) and squamous cell carcinoma data from (Puram et al., 2017) (2A-D). For both datasets, malignant cells were verified by bulk whole-exome sequencing. The aforementioned malignancy score accurately separated transformed, malignant cells producing areas under the receiver operating characteristic curve (AUROC) of 0.98 on the melanoma cells (Fig. 1E) and 0.97 on the squamous cell carcinoma cells (Fig. 2E).

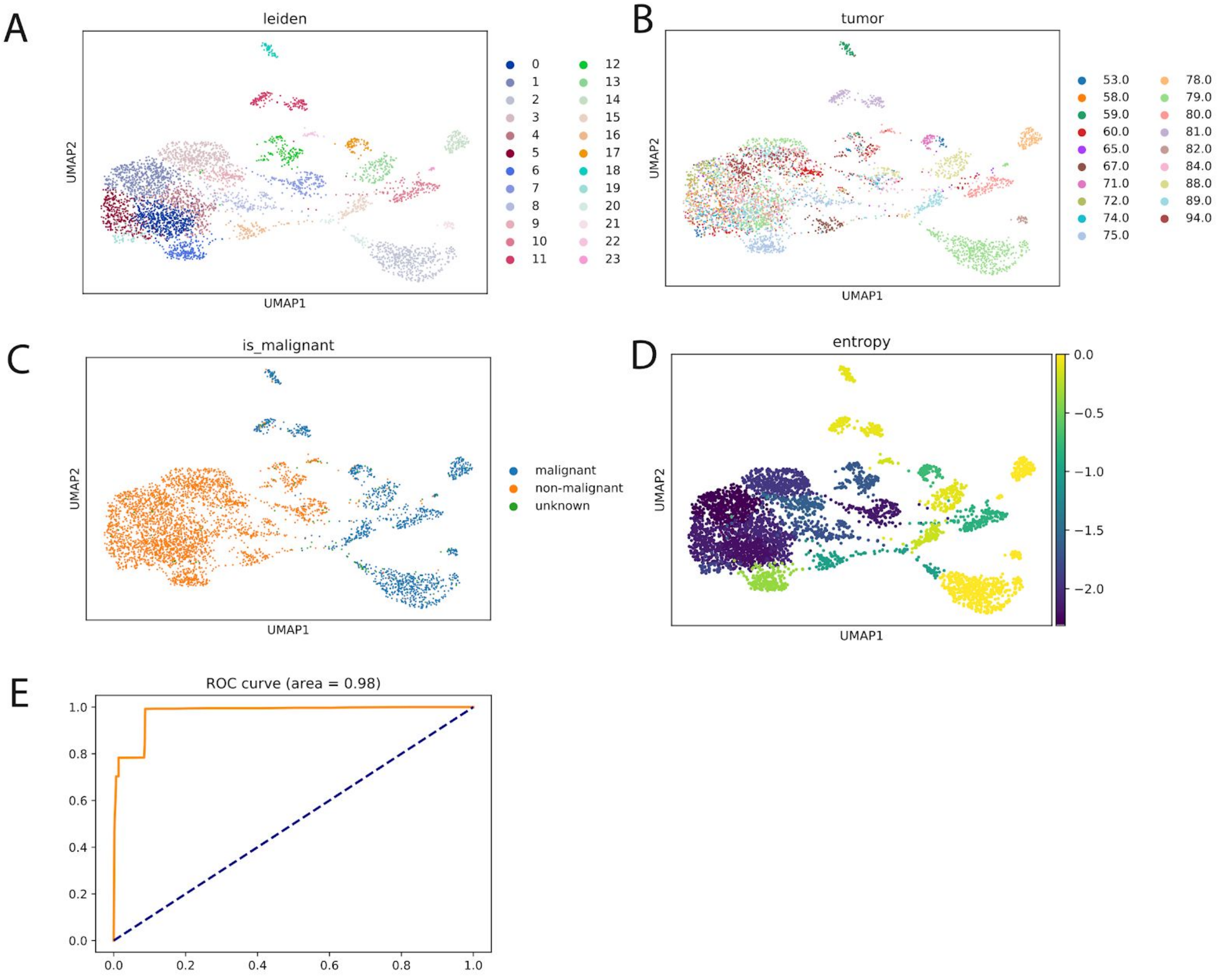


Figure 1. **Benchmarking the malignancy score on data from Tirosh et al. (2016).** UMAP plot of cells colored by cluster (**A**), tumor of origin (**B**), malignancy status as estimated by the authors of the dataset (**C**), and malignancy score (**D**). (**E**) The ROC curve evaluating the malignancy score using the malignancy status as estimated by the authors as the ground truth.

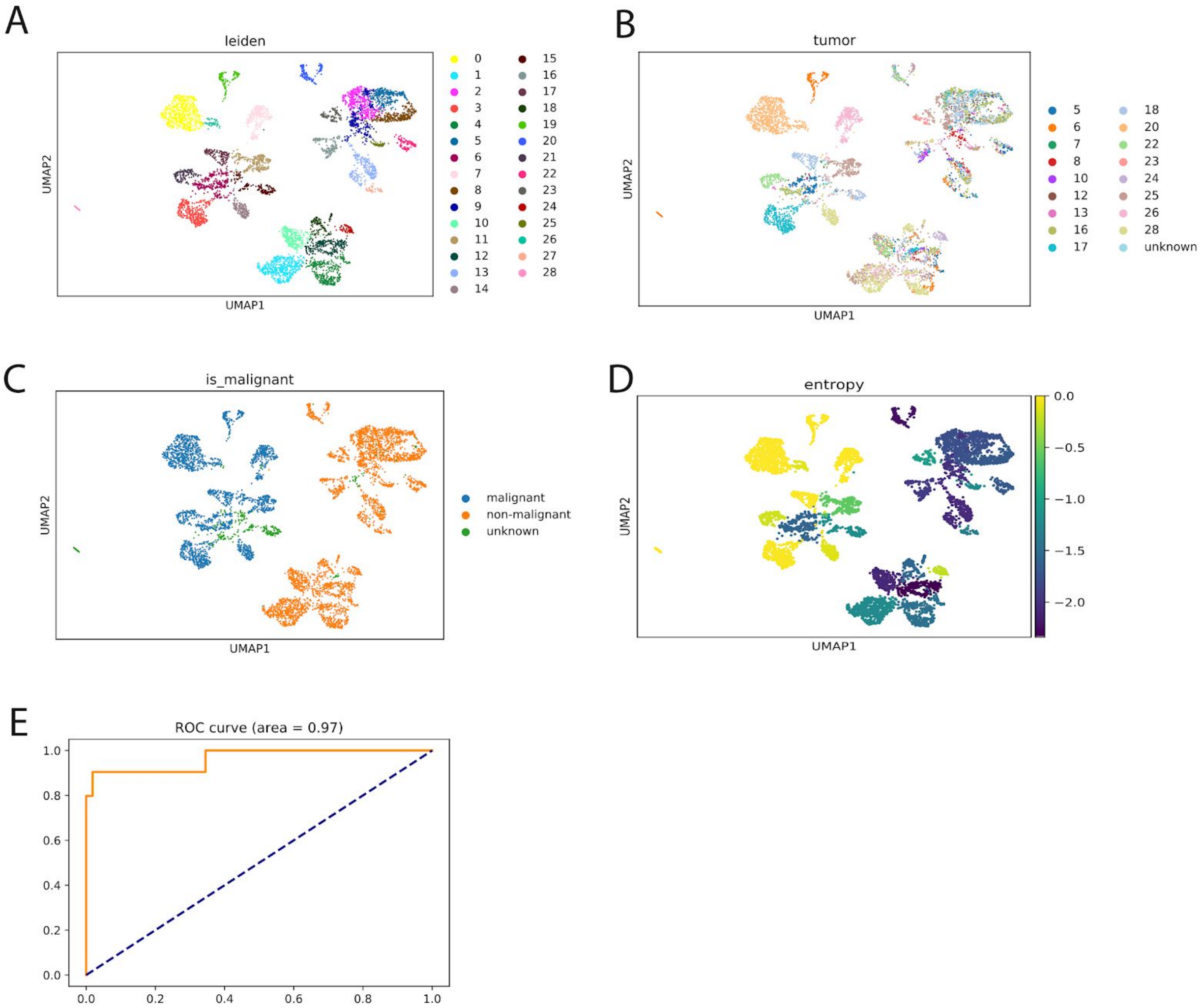


Figure 2. **Benchmarking the malignancy score on data from Puram et al. (2017).** UMAP plot of cells colored by cluster (**A**), tumor of origin (**B**), malignancy status as estimated by the authors of the dataset (**C**), and malignancy score (**D**). (**E**) The ROC curve evaluating the malignancy score using the malignancy status as estimated by the authors as the ground truth.

### Supplemental References


Couturier, C.P., Ayyadhury, S., Le, P.U., Nadaf, J., Monlong, J., Riva, G., Allache, R., Baig, S., Yan, X., Bourgey, M., et al. (2020). Single-cell RNA-seq reveals that glioblastoma recapitulates a normal neurodevelopmental hierarchy. Nat. Commun. *11*, 3406.

Puram, S.V., Tirosh, I., Parikh, A.S., Patel, A.P., Yizhak, K., Gillespie, S., Rodman, C., Luo, C.L., Mroz, E.A., Emerick, K.S., et al. (2017). Single-Cell Transcriptomic Analysis of Primary and Metastatic Tumor Ecosystems in Head and Neck Cancer. Cell *171*, 1611–1624.e24.

Tirosh, I., Izar, B., Prakadan, S.M., Wadsworth, M.H., 2nd, Treacy, D., Trombetta, J.J., Rotem, A., Rodman, C., Lian, C., Murphy, G., et al. (2016). Dissecting the multicellular ecosystem of metastatic melanoma by single-cell RNA-seq. Science *352*, 189–196.
